# Landscape-mediated spread dynamics and anticipatory surveillance of African swine fever in wild boar: insights from the 2025–2026 Catalonia outbreak in Spain

**DOI:** 10.64898/2026.07.03.736421

**Authors:** Jaime Bosch, Benjamin Ivorra, Cecilia Aguilar-Vega, Satoshi Ito, Jose Manuel Sánchez-Vizcaíno, Angel M. Ramos

## Abstract

African swine fever (ASF) in wild boar poses major surveillance and control challenges, particularly in periurban and human-modified landscapes where ecological complexity, delayed detection, and heterogeneous host movements complicate outbreak interpretation. In late 2025, ASF was detected in wild boar in Catalonia, Spain, creating a high-priority epidemiological scenario at the wildlife–urban interface. In this study, we analysed the 2025–2026 Catalonia outbreak using scenario-based spatial modelling with the WIMBOARD (Wild Integrated Movement Boar Outbreak and Risk Dynamics) framework. Simulations under baseline control conditions were used to interpret spread dynamics and support anticipatory surveillance. The results indicate that outbreak expansion was structured and directional rather than isotropic. Temporal outputs suggested delayed detectability between infection prevalence and mortality signals, whereas cumulative spatial risk maps, time-to-infection surfaces, and monthly infection-risk dynamics identified directional asymmetries and differentiated phases of spread. An early southward dispersal signal was consistent with initial field observations, while the north–northwest sector emerged as the most consequential expansion scenario because of its stronger functional connectivity and higher potential for regional amplification and persistence, while a later northeastward phase remained comparatively less influential within the 500-day simulation horizon. Monthly risk surfaces further showed that ASF spread behaved as a moving eco-epidemiological wavefront, allowing identification of shifting surveillance windows before mortality became apparent. These findings support the interpretation of the Catalonia outbreak as a landscape-mediated epidemiological process shaped by functional connectivity, wildlife behavioural adaptation, peri-urban ecological structure, and delayed detection. This study shows how spatially explicit eco-epidemiological modelling can support outbreak interpretation, surveillance prioritisation, and proactive wildlife disease management under complex field conditions.

## 1. Introduction

African swine fever (ASF) remains one of the most consequential transboundary animal diseases affecting both domestic pig production and wild boar populations worldwide (Costard et al., 2013). Characterised by complex transmission pathways, high environmental resistance and the capacity for both local persistence and long-distance dissemination, the disease has continued to expand geographically since the major Eurasian epidemic wave triggered by the 2007 introduction in Georgia (Dixon et al., 2020; Chenais et al., 2018). ASF has shown an unprecedented geographic expansion in recent decades, affecting multiple continents and generating complex wildlife-livestock epidemiological interfaces (Dixon et al., 2020). In Europe, its epidemiology has been increasingly shaped by the spatial ecology, demographic dynamics, and behavioural plasticity of wild boar, particularly in landscapes strongly modified by human activity (Morelle et al., 2019a; Bosch et al., 2012). In this context, wild boar populations have been recognised as a key wildlife reservoir contributing to the maintenance and spatial spread of ASF virus across heterogeneous landscapes (Bosch et al., 2017a,b; Chenais et al., 2018; Podgórski and Śmietanka, 2018; Pepin et al., 2020; Sauter-Louis et al., 2020). Under such conditions, the emergence and spread of ASF cannot be understood solely in terms of pathogen transmission, but must also be interpreted through the interaction among host ecology, landscape structure, and surveillance performance (Bosch and Ivorra et al., 2026a; Bosch et al., 2026b). The complexity of ASF epidemiology in Europe has been extensively documented, highlighting the role of wildlife reservoirs, surveillance performance and diagnostic capacity in shaping outbreak persistence and control effectiveness (Gallardo et al., 2015; Sánchez-Vizcaíno et al., 2012; Blome et al., 2020). ASF epidemiology in wild boar populations has also been recognised as strongly influenced by carcass-mediated environmental persistence, host movement ecology and landscape connectivity processes operating across heterogeneous ecological systems (Chenais et al., 2018; Morelle et al., 2019a; Bosch et al., 2017a; Goicolea et al., 2024; Faustini et al., 2025), with infected carcasses acting as a major driver of local virus maintenance and transmission risk in scenarios where high virulent strains circulate (Probst et al., 2017).

Against this broader European epidemiological background, the re-emergence of ASF in Spain (Catalonia) provides a particularly informative peri-urban case study illustrating how landscape structure, wildlife ecology and surveillance performance interact under real outbreak conditions (Bosch and Ivorra et al., 2026a. The confirmation of ASF in wild boar in Catalonia in late 2025 (World Organisation for Animal Health (WOAH), 2025) provides a relevant example of this epidemiological context, highlighting the challenges of interpreting and managing disease spread in heterogeneous and highly anthropised urban–natural interface environments. In this study, we refer to the affected populations as urban-adapted wild boar embedded within an urban–natural interface, reflecting their behavioural plasticity and ecological adaptation to human-modified environments. At the local scale, the outbreak is characterised by the particular epidemiological features of peri-urban ASF emergence in southern Europe. The first confirmed cases in late November 2025 were detected in wild boar carcasses in a highly anthropised landscape near the metropolitan area of Barcelona, specifically in the municipality of Cerdanyola del Vallès, close to the Autonomous University of Barcelona (UAB) and the Bellaterra residential area. The area is characterised by fragmented habitats, transport infrastructures and intense human activity, together with medium-sized forest patches connected to the Collserola Natural Park and adjacent natural areas. The absence of infected domestic pig farms at the time of detection and in subsequent observations, together with the presence of both fresh carcasses and skeletal remains, suggests that virus circulation may have preceded official notification, highlighting the challenge of delayed detectability in wildlife systems, where early infection phases are typically more difficult to identify than in managed domestic populations.

Spain is not unfamiliar with ASF epidemiology. The country experienced prolonged circulation of ASF virus genotype I from the 1960s until its official eradication in 1995, representing one of the most extensive control programmes against the disease in Europe (Arias and Sánchez-Vizcaíno, 2002; Sánchez-Vizcaíno et al., 2012). This historical experience has contributed to the development of strong institutional, technical and operational capacity, including long-standing surveillance systems in both domestic pigs and wild boar, contingency planning, and high biosecurity standards in commercial production systems. Previous work has highlighted the importance of such preparedness frameworks in enabling rapid detection and response under ASF risk scenarios (Mur et al., 2012; Sánchez-Vizcaíno et al., 2015). However, despite this accumulated expertise, ASF epidemiology remains characterised by unpredictable long-distance introductions, often associated with human-mediated pathways such as contaminated pork products or fomites (Costard et al., 2013; Chenais et al., 2018; Aguilar-Vega et al., 2024). In this context, preparedness does not necessarily prevent introduction, but rather determines the capacity to respond effectively once the virus is detected. The reemergence of ASF in Catalonia therefore occurs within a setting combining historical epidemiological experience, high preparedness, and persistent vulnerability to stochastic introduction events. From a broader historical perspective, the epidemiology of ASF in the Iberian Peninsula was also influenced by ecological mechanisms beyond direct host-to-host transmission. In particular, soft ticks of the *Ornithodoros erraticus* complex acted as long-term biological vectors and virus reservoirs in some extensive pig-production systems in southwestern and central Spain (Boinas et al., 2011; Sánchez-Vizcaíno et al., 2012; Boinas et al., 2014). This historical context should be taken into account when interpreting ASF epidemiology in Spain, as tick-mediated transmission played an important role during the earlier Spanish epidemic. However, in the Catalonia setting, based on the information currently available, the presence of this vector has not been reported in the affected region based on currently available distribution data (GBIF, 2023). Its role can therefore be considered negligible in the present outbreak scenario.

Control of ASF in wild boar populations has often relied on reactive measures implemented after outbreak confirmation, including passive surveillance, carcass removal, movement restrictions, and population management interventions. Early detection of infection and rapid implementation of coordinated control measures have been repeatedly identified as critical determinants of successful ASF containment strategies in both domestic pigs and wild boar populations (Costard et al., 2013; Gallardo et al., 2015; Sánchez-Vizcaíno et al., 2015). Population-management interventions such as hunting pressure modulation or abundance reduction have shown variable epidemiological effectiveness depending on landscape structure and host behavioural responses (Keuling et al., 2013; Pepin et al., 2020). Modelling studies have further suggested that spatial coordination and timing of host reduction measures are critical for limiting epidemic expansion in wildlife systems (Lange and Thulke, 2017). Although these actions remain essential components of ASF control, their effectiveness may be limited by delayed detection, uncertainty regarding underlying infection dynamics, and the complex interaction between ecological and anthropogenic processes affecting host movement and contact patterns (Bosch et al., 2012; Guinat et al., 2016; Chenais et al., 2018). In fragmented periurban-natural systems, where habitat discontinuities, anthropogenic resources, and residual connectivity structures coexist, these limitations may be even more pronounced. This raises a central operational challenge: how can surveillance and control be informed before visible mortality patterns emerge and infection becomes widely established across the landscape?

Recent advances in experimental epidemiology have improved our understanding of ASF transmission in wild boar, including the possible influence of virulence variation on host survival, infectious-period duration, and outbreak persistence (Martínez & Bosch et al., 2023). At the same time, landscape-based epidemiological approaches have shown that wild boar movement and disease spread are strongly conditioned by habitat suitability, ecological corridors, and barriers to functional connectivity (Morelle and Lejeune, 2015; Bosch et al., 2017a; Podgórski et al., 2018; Goicolea et al., 2024). The development of such operational modelling platforms increasingly relies on multidisciplinary and interdisciplinary collaboration, combining expertise in epidemiology, ecology, veterinary science, environmental analysis and applied mathematics to address the inherently complex nature of wildlife disease dynamics under real-world conditions.

In this study, we analyse the 2025–2026 ASF outbreak in wild boar in Catalonia from an eco-epidemiological and spatial-risk perspective. Using scenario-based modelling within the WIMBOARD framework (Wild Integrated Movement Boar Outbreak and Risk Dynamics), developed within the framework (European Commission and VACDIVA Consortium, 2024), we examine temporal epidemic dynamics, cumulative spatial risk patterns, directional dispersal phases, and monthly infection-risk surfaces in order to interpret likely spread trajectories and their operational implications. Additional field-operational and preliminary spatial-epidemiological insights from the evolving scenario have been reported elsewhere (Bosch and Ivorra et al., 2026a; Bosch et al., 2026b). The analytical framework used in this study builds on previous experience in spatial and stochastic epidemiological modelling developed for livestock diseases, including earlier modelling systems such as Be-FAST for classical swine fever and related disease-spread applications (Martínez-López et al., 2011, 2012; Ivorra et al., 2014; Fernández-Carrión et al., 2016). Within this broader modelling trajectory, WIMBOARD was developed as a wildlife-oriented framework specifically designed to analyse ASF spread in wild boar populations under heterogeneous landscape conditions and control measures. Its development was grounded in a strongly interdisciplinary philosophy, bringing together applied mathematics, epidemiology, veterinary science, wildlife biology, ecology, environmental processes and landscape connectivity analysis in order to address the multidimensional nature of ASF spread in wild boar systems. Before its application to the Catalonia outbreak, WIMBOARD had been progressively evaluated in other European ASF contexts, including outbreak scenarios in northern Italy and Belgium during 2022–2023, where its spatiotemporal projections were contrasted with field epidemiological conditions. This prior translational experience supports the operational maturity of the framework and helps position the present Catalonia application not as an isolated modelling exercise, but as the adaptation of an already functional eco-epidemiological platform to an emerging ASF scenario in Spain. Its transfer-oriented character was also reflected at the VACDIVA final technology-transfer workshop, held at the European Economic and Social Committee (EESC) in Brussels in November 2024, where WIMBOARD was presented as a European outcome for ASF preparedness and control. Rather than focusing on model development itself, this study uses spatially explicit modelling as a tool to investigate how functional landscape connectivity, periurban ecological structure, and delayed detectability may shape ASF spread at a complex wildlife–urban interface. Operating seamlessly from continental-scale European epidemiological modelling to fine-scale local risk resolution, WIMBOARD converts large-scale ecological complexity into actionable spatial intelligence for precision surveillance and adaptive outbreak control.

The objectives of this study are threefold: first, to interpret the 2025-2026 Catalonia outbreak as an eco-epidemiological process structured by landscape and host ecology; second, to identify spatial and temporal features of outbreak expansion that are relevant for anticipatory surveillance and containment; and third, to illustrate how spatially explicit scenario analysis can support field interpretation and decision-making in complex wildlife disease systems. In doing so, the study aims to contribute to a more proactive and operationally relevant understanding of ASF dynamics in peri-urban wild boar populations.

The remaining of the paper is organised as follows. Section 2 presents the epidemiological context of ASF emergence in Catalonia, including the outbreak setting, the main sources of uncertainty, and the relevance of the peri-urban wildlife–urban interface. Section 3 describes the analytical framework and scenario-based modelling approach used in this study. Section 4 presents the main results and their eco-epidemiological interpretation, with emphasis on temporal dynamics, spatial risk structure, directional dispersal, and operational surveillance implications. This study also introduces and operationalises the concept of landscape-mediated epidemiological modulation to explain how fragmented urban-agro-natural mosaics can alter transmission structure, mortality visibility and surveillance performance. Finally, the paper closes with the main conclusions and broader implications for anticipatory wildlife disease management.

## 2. Epidemiological context of ASF emergence in Catalonia

The re-emergence of ASF in Spain in late November 2025 marked a major epidemiological event for the Iberian Peninsula after more than three decades without confirmed virus circulation. The first detected cases were reported between 25 and 26 November 2025, when two wild boar carcasses were found in the municipality of Cerdanyola del Vallès, within the metropolitan area of Barcelona in northeastern Spain (World Organisation for Animal Health (WOAH), 2025). The initial outbreak focus was located in a highly anthropised peri-urban landscape characterised by a mosaic of residential areas, transport infrastructures, and fragmented forest and agroforestry patches connected to the Collserola Natural Park. Wild boar abundance in this area has been considered medium to high, reflecting the species’ increasing adaptation to human-dominated environments (Alexander et al., 2016; Pittiglio et al., 2018).

### 2.1. Outbreak setting and initial response

From the onset, the outbreak displayed several features of particular epidemiological relevance. Detection was initially limited to carcasses, with no infected live animals or affected domestic pig farms identified. Necropsy findings and laboratory confirmation by the national reference laboratory established the presence of ASF virus genotype II, confirming the disease in Spain for the first time since the official eradication of genotype I in 1995 (Arias and Sánchez-Vizcaíno, 2002). The later detection of skeletal remains suggested that viral circulation may have preceded official notification by several weeks or months, consistent with the delayed detectability often observed in wildlife disease outbreaks. The 2025-2026 Catalonia outbreak therefore represents not only a sanitary emergency, but also a real-time epidemiological situation in which preparedness plans, wildlife management strategies, and surveillance systems are being tested simultaneously under field conditions.

Following confirmation, a coordinated emergency response was activated. Measures included the establishment of a high-risk infection core zone with a radius of 6 km and a surrounding surveillance buffer zone extending up to 20 km. Intensive carcass search and recovery operations were implemented using trained personnel, detection dogs, aerial surveillance, and drone technology. Traditional hunting activities were temporarily suspended to minimise disturbance-driven dispersal of wild boar, while targeted population management based on trapping and controlled removal was applied to reduce host abundance. Professional selective culling was also implemented by trained personnel, in contrast to traditional hunting, supported by the use of suppressed firearms to minimise disturbance and avoid unintended dispersal effects. Reinforcement of farm biosecurity and restrictions on human activities in the natural environment were also prioritised in order to reduce the risk of indirect virus spread. These interventions were broadly consistent with established European standards for ASF control in wildlife and reflected the high level of preparedness developed in Spain through surveillance programmes and contingency planning (European Commission, 2020; Gervasi et al., 2022; European Commission, 2023).

Despite rapid operational deployment, several epidemiological uncertainties remained. As of early 2026, all confirmed cases were still confined to wild boar populations, and spatial containment initially appeared to be effective. However, the outbreak occurred in an ecological setting markedly different from many previous European ASF events, which were often detected in more remote and less anthropised landscapes. In periurban-natural ecosystems such as Collserola, indirect human-mediated transmission pathways, including contaminated fomites, waste accessibility, and intense recreational use of natural areas, may become as relevant as natural host movement in shaping spatial transmission dynamics (Bosch et al., 2017b; De la Torre et al., 2015). This makes outbreak interpretation particularly challenging and strengthens the need for analytical approaches capable of integrating landscape structure, host ecology, and surveillance constraints.

### 2.2. Molecular and epidemiological uncertainty

From a molecular epidemiological perspective, the available public information indicates that the detected virus belongs to genotype II, associated with the Eurasian epidemic lineage since 2007, but presents distinctive genomic characteristics that differentiate it from currently sequenced circulating European isolates (group 2-28). The strain has been provisionally classified as belonging to a new genetic group (Group 29), suggesting an unexpectedly high level of divergence for a large double-stranded DNA virus with a relatively low evolutionary rate (Gámbaro et al., 2025; Nefedeva et al., 2020). On the basis of currently available information, the precise origin of the outbreak cannot be confirmed. Hypotheses under consideration include sporadic long-distance introduction linked to human activities, undetected circulation in wildlife, and other scenarios requiring additional genomic evidence. However, as in previous European outbreaks, clarifying the origin remains secondary to the immediate priority of interrupting transmission and containing spatial spread.

Additional working hypotheses include sporadic introduction associated with human-mediated activities, such as the disposal of contaminated pork products or waste accessible to wild boar. Initially, an accidental release from facilities handling ASF virus or experimental vaccines was also considered but rejected with full-genome sequencing. However, the corresponding phylogenetic trees and detailed sequencing data have not yet been publicly released, and therefore independent evaluation is currently limited. Intentional introduction cannot be formally excluded at this stage. Conversely, progressive natural spread through wild boar movements from the nearest known infected areas in Europe appears epidemiologically less likely, given the species’ limited long-distance dispersal capacity and the absence of spatial epidemiological continuity across intervening regions. In the absence of direct evidence, most introduction pathways remain under investigation.

Another critical dimension concerns the apparent epidemiological behaviour of the circulating strain. Field observations may suggest moderate mortality patterns that have prompted consideration of a possible lower-virulence scenario. Early detection of seropositive wild boar during the initial outbreak phase may indicate prolonged host survival, potentially associated with altered virulence expression or heterogeneous infection outcomes. In wildlife populations, moderately virulent or attenuated ASF strains may generate more complex transmission dynamics than highly virulent variants. Increased survival time of infected individuals may allow longer dispersal distances, more diffuse spatial clustering, and lower detectability through passive surveillance based on carcass recovery. These mechanisms may complicate the delimitation of infected areas and prolong environmental virus persistence. Experimental evidence derived from controlled infection studies in wild boar has shown that coexistence scenarios involving strains of different virulence may produce heterogeneous transmission patterns and intermittent persistence dynamics (Martínez & Bosch et al., 2023). Although the epidemiological characteristics of the 2025-2026 Catalonia outbreak strain remain under investigation, this uncertainty reinforces the need for adaptive surveillance strategies capable of incorporating multiple plausible transmission scenarios.

### 2.3. Peri-urban ecological context and analytical implications

Beyond virological and operational uncertainty, the outbreak also reflects a broader ecological reality. Wild boar populations across Europe have undergone major demographic and behavioural changes driven by anthropogenic landscape transformation (Barrios-García and Ballari, 2012; Bosch et al., 2012). As opportunistic omnivores, wild boar exploit a broad range of natural and anthropogenic food resources in heterogeneous landscapes. Expansion of urbanagro–forest interfaces, year-round food availability associated with agricultural intensification, the absence of large predators, and historically uncoordinated hunting practices have all contributed to increased abundance and altered movement ecology (Amici et al., 2012; Morelle and Lejeune, 2015; Bosch et al., 2017a; Aguilar-Vega et al., 2024). As opportunistic scavengers capable of consuming carcasses and anthropogenic waste, wild boar are also particularly exposed to environmentally persistent pathogens such as ASF virus. In this context, the species should not be regarded solely as a disease environmental source, but also as an ecological amplifier linking environmental disturbance, host demography, and pathogen transmission within human-modified landscapes.

Overall, the 2025-2026 Catalonia outbreak illustrates the convergence of molecular uncertainty, ecological complexity, and operational challenges that characterise transboundary wildlife diseases in urban–natural interface settings. It also highlights the limitations of purely reactive response frameworks. Analytical tools capable of integrating landscape connectivity, host movement, surveillance performance, and alternative epidemiological scenarios may therefore play an important role in supporting outbreak interpretation, surveillance prioritisation, and preparedness planning in such dynamic contexts. Such integrative approaches may also facilitate earlier detection of secondary dispersal fronts in neighbouring or previously unaffected areas. To analyse this emerging epidemiological configuration more robustly, the following section presents the analytical framework and scenario-based modelling approach used to interpret outbreak dynamics and explore plausible spread scenarios under the Catalonia setting.

## 3. Analytical framework and scenario-based modelling approach

This study combines experimental epidemiological evidence (Martínez & Bosch et al., 2023), field interpretation, and scenario-based spatial modelling to analyse the 2025–2026 ASF outbreak in wild boar in Catalonia. The analytical framework was designed to address a central difficulty of this outbreak: the need to interpret disease spread under conditions of ecological complexity, delayed detectability, and epidemiological uncertainty in a fragmented peri-urban landscape with interconnected natural habitat patches. Rather than relying exclusively on reactive outbreak indicators, the approach integrates biological knowledge on ASF transmission with spatially explicit modelling in order to explore plausible spread trajectories and their implications for surveillance and control.

### 3.1. Epidemiological rationale for scenario-based analysis

Understanding ASF dynamics in wild boar populations requires combining insights from theoretical transmission modelling and experimental infection studies. Classical epidemiological frameworks have long emphasised the role of virulence gradients in shaping outbreak dynamics, particularly through their influence on infectious-period duration, mortality, and long-term persistence within host populations. Under highly virulent scenarios, outbreaks are typically characterised by rapid disease progression, acute mortality, and relatively short infectious periods, often generating spatially clustered events that may be more readily detected through passive carcass-based surveillance (Blome et al., 2020; Probst et al., 2017; Costard et al., 2013). By contrast, lower-virulence or moderately pathogenic scenarios may involve prolonged host survival, reduced lethality, and more diffuse spatial transmission patterns, thereby increasing the risk of delayed detection and extended environmental contamination (Martínez & Bosch et al., 2023).

Experimental epidemiology provides an essential complement to these theoretical expectations by quantifying infection outcomes under controlled conditions. Recent infection studies in wild boar have shown that ASF transmission dynamics may be more heterogeneous than those represented in simplified deterministic models. In particular, coexistence scenarios involving strains with different virulence profiles may generate variable mortality trajectories, prolonged survival of seropositive individuals, and intermittent transmission windows extending beyond the acute outbreak phase (Martínez & Bosch et al., 2023). These findings suggest that field epidemiological signals may reflect a composite interaction between pathogen virulence, host demography, behavioural ecology, and environmental persistence, rather than a single dominant transmission regime.

This perspective is especially relevant for the Catalonia outbreak due to the possible circulation of a less virulent strain. Preliminary field observations, including the detection of seropositive wild boar and reports of heterogeneous mortality timing, suggest that the observed epidemiological pattern may not fully conform to a strictly classical acute high-virulence genotype II scenario. Under such conditions, carcass-based surveillance may underestimate the true extent and duration of viral circulation, hindering control strategies. These uncertainties justify the use of a scenario-based analytical framework capable of integrating multiple biologically plausible transmission configurations.

### 3.2. WIMBOARD as a scenario-based modelling framework

To support interpretation of the Catalonia outbreak, we used the Wild Integrated Movement Boar Outbreak and Risk Dynamics (WIMBOARD) framework as a scenario-based eco-epidemiological modelling tool (European Commission and VACDIVA Consortium, 2024). The analytical approach employed here builds on previous experience in spatial and stochastic epidemiological modelling for livestock diseases, including earlier systems such as Be-FAST for classical swine fever and related disease-spread applications (Martínez-López et al., 2011, 2012; Ivorra et al., 2014; Fernández-Carrión et al., 2016). Within this broader modelling trajectory, WIMBOARD was developed as a wildlife-oriented framework designed to analyse ASF spread in wild boar populations under heterogeneous landscape conditions. In the present study, it was used to explore plausible ASF spread trajectories under the Catalonia setting rather than as the primary object of methodological investigation. The first ASF detections in Catalonia were reported on 25–26 November 2025. Between 26 and 28 November, the outbreak scenario was prepared using all publicly available and verified information. By 1 December 2025, predictive model outputs were already available to support early epidemiological interpretation and response planning.

At an operational level, the framework integrates four main components or modules: (i) a spatial movement component informed by wild boar ecology and landscape structure; (ii) a population-dynamics component varying across space and time; (iii) an epidemiological transmission component representing infection progression under ASF outbreak conditions, including exposed (E), infectious (I), dead (D), recovered (R), and vaccinated (V) compartments, although the vaccination component was not activated in the present application; and (iv) configurable intervention modules allowing the exploration of control and surveillance scenarios. The WIMBOARD platform incorporates a compartmental state-transition transmission model of SEIR type coupled to a spatially explicit wild boar movement layer informed by ecology, landscape connectivity, and configurable control interventions. Through this structure, the framework links host movement, functional connectivity, epidemiological progression, and management actions within a single spatially explicit analytical environment.

This approach is particularly relevant in peri-urban systems, where habitat fragmentation, anthropogenic resources, movement barriers, and residual ecological corridors may strongly modulate transmission dynamics. In this context, scenario-based modelling is not intended to predict outbreak evolution deterministically, but to identify ecologically plausible directions of spread, zones of amplification, temporal surveillance windows, and areas in which delayed visibility may compromise outbreak interpretation.

### 3.3. Application to the 2025-2026 Catalonia outbreak setting

For the Catalonia application, the model was initialised on 1 November 2025 using the location of the first officially detected ASF-positive wild boar case and assuming, on the basis of field epidemiology and wild boar ecology, that viral circulation had probably begun before official notification. This assumption was consistent with the later detection of ASF-positive skeletal remains, suggesting that the outbreak may already have been under way before the first confirmed report. All simulations were conducted under baseline control conditions designed to reflect, as closely as possible, the control measures implemented in the real outbreak setting.

The modelling application was designed to represent the ecological and epidemiological characteristics of the affected peri-urban landscape, including fragmented habitat structure, wild boar presence in the Barcelona–Collserola–Vallès system, and the spatial heterogeneity associated with urban–natural interfaces, transport infrastructures, agroforestry mosaics, and residual connectivity corridors. Within this setting, the model was used to explore short- and medium-term spread scenarios under a plausible period of pre-detection transmission and under epidemiological assumptions broadly consistent with ASF circulation in wild boar.

Importantly, the analytical objective was not to reconstruct the outbreak retrospectively in a purely descriptive way, but to generate spatially and temporally structured scenarios that could help interpret the evolving epidemiological situation. In this sense, the Catalonia application functioned as a real-time modelling exercise supporting field interpretation under uncertainty.

### 3.4. Outputs used for outbreak interpretation

The analysis focused on a set of outputs selected for their epidemiological and operational relevance. These included temporal epidemic curves, prevalence dynamics, cumulative spatial risk maps, time-to-infection surfaces, and monthly infection-risk maps over the simulation horizon. Together, these outputs provided complementary information on outbreak timing, likely spread directions, spatial amplification patterns, and the possible mismatch between infection circulation and visible mortality.

Temporal outputs were used to assess the relationship between infection prevalence, mortality signals, and delayed detectability. Cumulative spatial risk maps and time-to-infection surfaces were used to evaluate the structure of spread in relation to landscape connectivity and directional asymmetries. Monthly infection-risk maps were used to identify moving wavefronts of infection and to infer shifting surveillance windows in space and time. Taken together, these outputs supported an eco-epidemiological interpretation of the 2025-2026 Catalonia outbreak in which spread is treated as a landscape-mediated process rather than as a homogeneous radial expansion.

The following section presents the interpretation of these temporal and spatial outputs, with particular attention to delayed detectability, functional connectivity, peri-urban ecological modulation, and their implications for anticipatory surveillance and outbreak management.

## 4. Results and eco-epidemiological interpretation

This section presents the main outputs generated for The WIMBOARD 2025–2026 Catalonia outbreak scenario and interprets them from an eco-epidemiological perspective. Rather than treating these outputs as purely mathematical projections, we analyse them as the epidemiological expression of interacting ecological, behavioural, landscape, and anthropogenic processes shaping ASF spread in a fragmented peri-urban wild boar system. Taken together, the results indicate a structured, directional, and landscape-mediated epidemic rather than a homogeneous radial spread, with direct implications for surveillance and outbreak management.

### 4.1. Temporal dynamics and delayed detectability

The simulated epidemiological and prevalence trajectories (Fig. 1) indicate a system initially dominated by exposed (E), infectious (I), and dead compartments (D), while recovery (R), and the associated temporary immunity, remain limited during the early and intermediate phases of the outbreak. The combined exposed-plus-infectious prevalence remains epidemiologically dominant throughout most of the simulation, whereas immune prevalence increases only gradually and remains substantially lower than active infection prevalence. Operationally, this is important because the visible epidemic signal, especially under passive carcass-based surveillance, may lag behind the true infection front.

**Figure 1:**
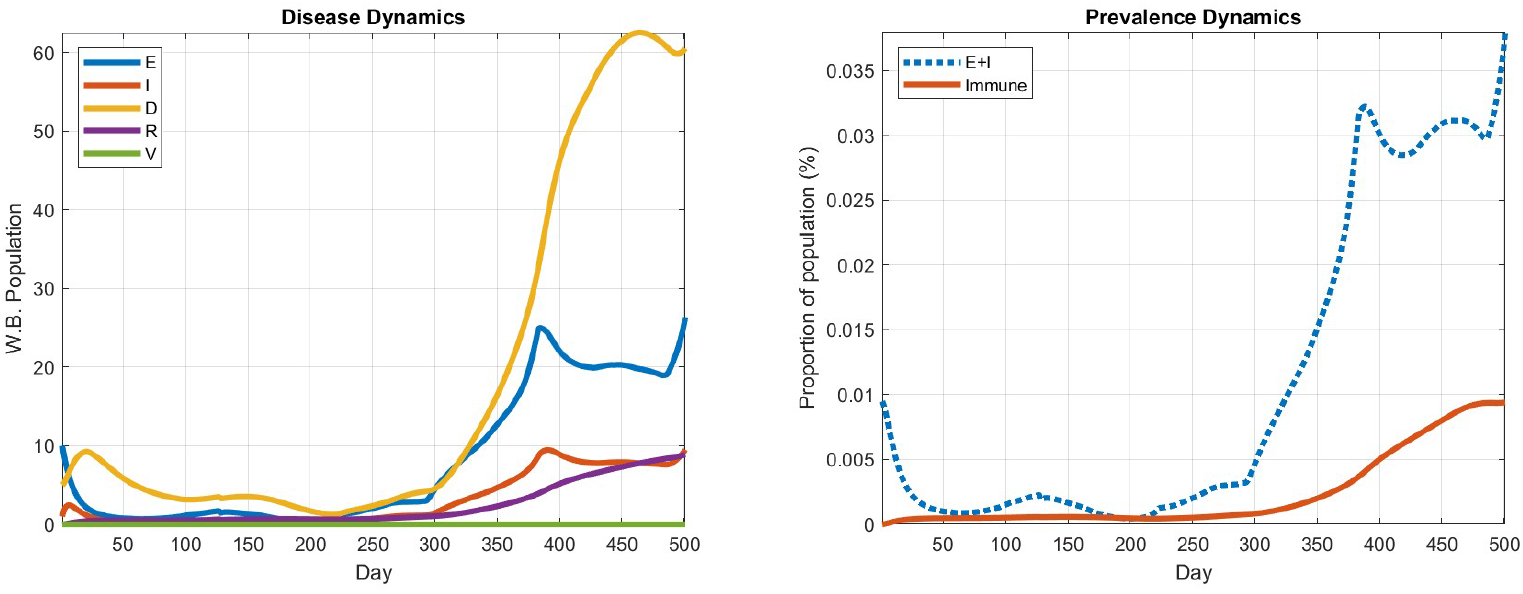
Simulated epidemiological and prevalence dynamics of African swine fever (ASF) in wild boar populations under the 2025–2026 Catalonia outbreak scenario over a 500-day simulation period. **Left:** simulated disease dynamics showing the temporal evolution of the exposed (E), infectious (I), dead (D), recovered (R), and vaccinated (V) compartments (not considered here). The modelled dynamics indicate delayed mortality relative to infection progression, with mortality becoming dominant during the later stages of the simulation and recovery increasing progressively towards the end of the time horizon. **Right:** simulated prevalence dynamics showing the temporal evolution of exposed-plus-infectious prevalence (E+I) in comparison with immune prevalence (R). The results highlight low early detectability followed by a progressive escalation of infection pressure, while immune prevalence increases gradually but remains lower than active infection prevalence throughout the simulated period.

A more detailed examination of the temporal trajectories shows a structured epidemic progression. The exposed compartment remains low during an apparent latency period and then increases sharply after approximately day 300, reaching a first major peak around days 380–400. When interpreted jointly with the spatial-risk outputs, this temporal escalation phase coincides with the emergence of structured northern expansion patterns, highlighting how the integration of temporal epidemic dynamics and spatial connectivity modelling improves the eco-epidemiological interpretability of outbreak spread. The infectious compartment follows shortly afterwards, reaching its maximum around days 390–410. Mortality increases later, rising steeply between approximately days 360 and 450 and becoming the dominant compartment towards the end of the simulation period. This delayed mortality signal suggests that a substantial fraction of infected animals remains mobile and epidemiologically active for a considerable period before death.

Unlike the initial version of the scenario, the updated trajectories also show a progressive increase in the recovered compartment during the later stages of the simulation. This is reflected in the prevalence panel, where immune prevalence rises steadily after approximately day 350, although it remains clearly below the exposed-plus-infectious prevalence throughout the simulation horizon. Thus, while the outbreak dynamics remain largely driven by ongoing transmission and mortality, the results also suggest a gradual accumulation of post-infection immune individuals in the population.

These temporal epidemiological patterns, characterized by delayed mortality signals, sustained active transmission and the gradual accumulation of post-infection immune individuals, are ecologically plausible in the fragmented urban–agro–natural landscape of the outbreak area, where wild boar populations exhibit modified movement routines, reduced predictability of long-range dispersal, and contact networks shaped by anthropogenic resources and habitat discontinuities. Under such conditions, infection may spread progressively while remaining only partially visible to surveillance. Consequently, passive surveillance based primarily on carcass detection may underestimate the true spatial extent of viral circulation, and areas that appear epidemiologically quiet may already be undergoing active transmission. The Catalonia outbreak therefore supports the interpretation of ASF spread as a case of delayed detectability and landscape-mediated epidemiological modulation in a peri-urban wildlife system.

### 4.2. Spatial risk structure and directional dispersal

The cumulative spatial risk output at 500 days (Fig. 2) and the corresponding time-since-infection surface (Fig. 3) show that spread is neither isotropic nor simply distance-dependent. Instead, the simulated pattern displays clear directional asymmetries and differentiated phases of expansion, indicating that outbreak spread is being shaped by functional connectivity, habitat structure, barriers, corridors, and host ecology rather than by homogeneous diffusion. In this sense, the analysis reconstructs a biologically plausible geography of transmission that is consistent with connectivity-driven spread (Bosch et al., 2017a; Goicolea et al., 2024; Connectivity Modelling Project, 2024). All ASF cases in wild boar shown in the map were updated up to 17 March 2026 according to WOAH notifications.

**Figure 2:**
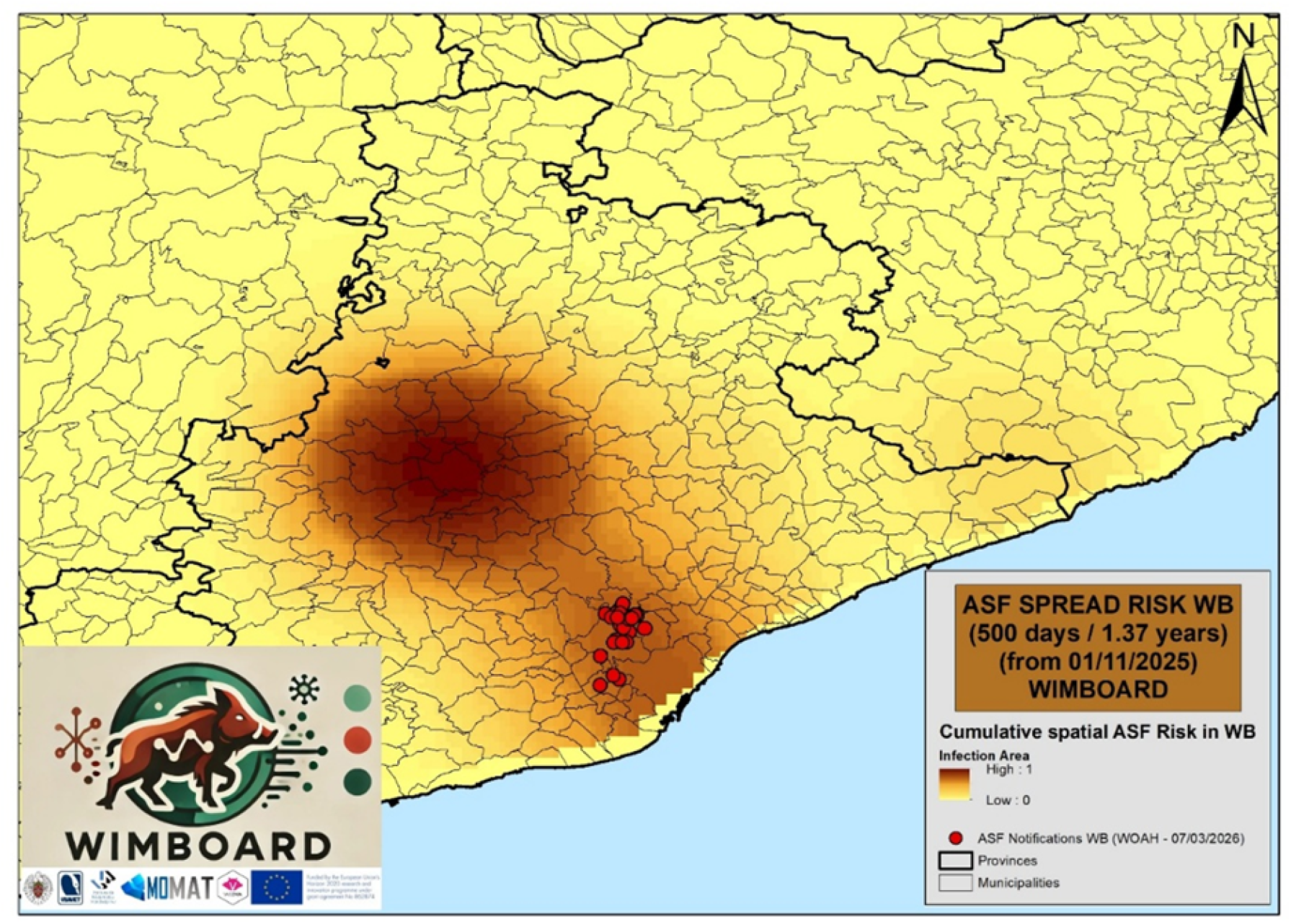
Cumulative spatial risk of African swine fever (ASF) infection after 500 days of simulated spread. Spatial distribution of normalised infection risk across the landscape, representing the integrated epidemiological pressure accumulated during approximately 1.37 years following the first notified infection. The spatial pattern reveals directional spread driven by functional connectivity rather than isotropic diffusion. ASF notifications in wild boar updated as of 17 March 2026, source: WOAH.

**Figure 3:**
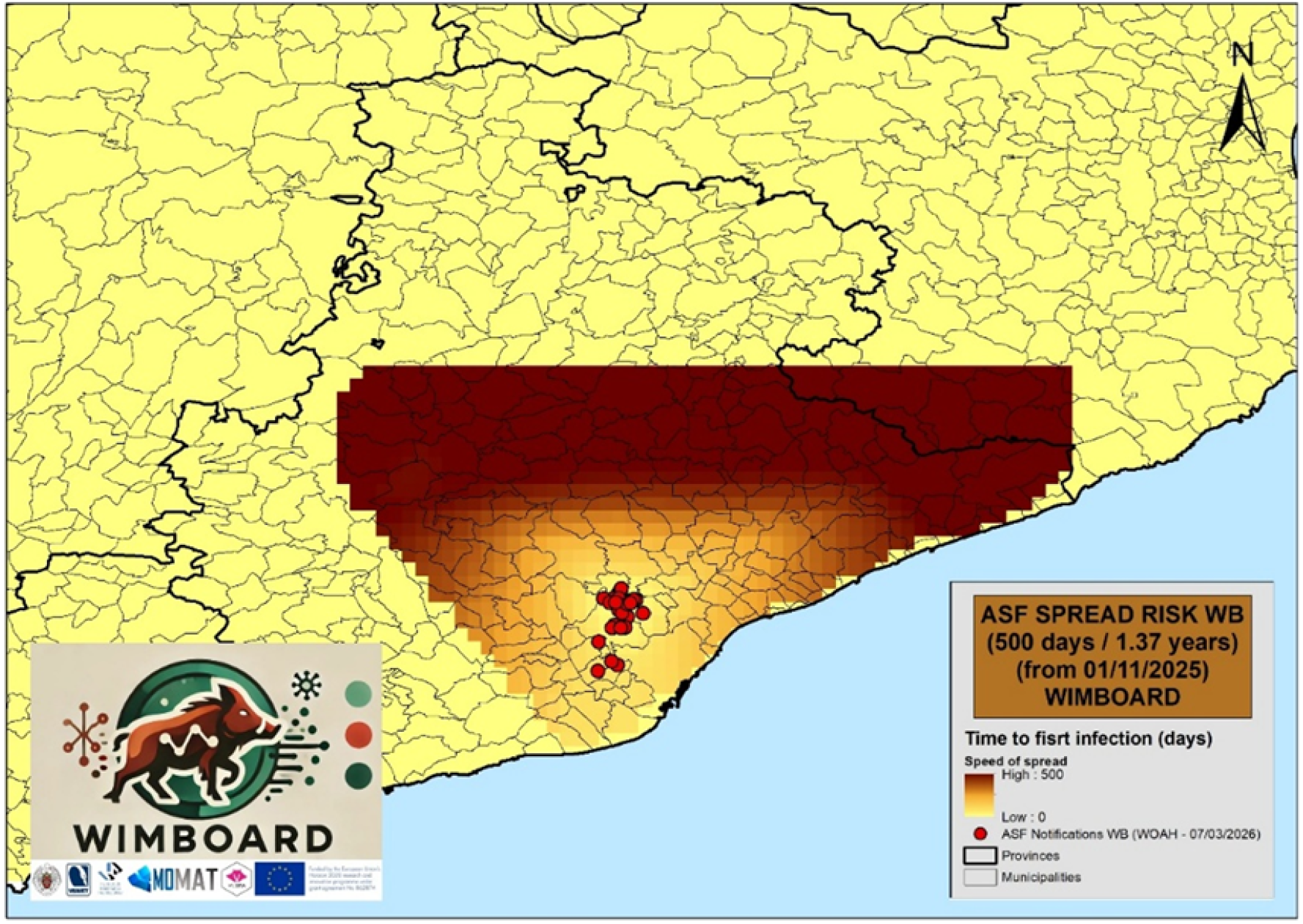
Time-since-infection surface after 500 days of simulated African swine fever (ASF) spread. Spatial distribution of the estimated timing of infection across the landscape, showing differentiated phases of epidemic expansion from the initial notified focus. The resulting pattern highlights marked directional asymmetries and supports the interpretation of a landscape-mediated transmission process structured by functional connectivity rather than by homogeneous distance-based diffusion.

Over the 500-day horizon, three main dispersal phases emerge. The first is an early southward phase, extending approximately from days 30 to 90 towards the Collserola system and the peri-urban southern sector. The second, between approximately days 250 and 350, is oriented towards the north and northwest and represents the phase of greatest infection pressure and cumulative epidemiological impact. The third, between approximately days 400 and 490, is directed towards the northeast and appears less intense, although it still reflects continued spatial progression. This third phase could become increasingly important beyond the 500-day simulation horizon; however, such long-term projections involve substantially greater uncertainty regarding their correspondence with the real outbreak trajectory. These phases are ecologically interpretable as successive expressions of how wild boar movement, local transmission, environmental contamination, and landscape structure interact over time.

The second phase is especially important because it points to a possible epidemiological amplification landscape within 500 simulation days. Compared with the initial southward signal, the northern sector is characterised by higher habitat quality, greater population abundance, and stronger functional connectivity at broader spatial scales (Bosch et al., 2017a; Goicolea et al., 2024; Connectivity Modelling Project, 2024). Under these conditions, infection spread could shift from a structured wave-front process to a broader regional expansion dynamic, increasing persistence, spatial unpredictability, and long-term control difficulty. The northern sector may therefore be interpreted as a zone with increased potential for multidirectional spread and long-term persistence. By contrast, the later northeastward phase appears to reflect a more attenuated secondary expansion front advancing through progressively connected habitat elements. Although its projected epidemiological impact is lower, it still highlights the need for sustained surveillance beyond the initial epidemic wave.

### 4.3. Peri-urban modulation of spread and monthly wavefront dynamics

When cumulative spatial risk (Fig. 2), time-to-infection gradients (Fig. 3), and the monthly infection-risk sequence (Fig. 4) are considered together, they support the interpretation of ASF spread as a landscape-mediated process in a fragmented peri-urban system. The cumulative map captures the long-term geography of epidemiological pressure, whereas the monthly surfaces reveal the progressive and wave-like nature of the epidemic front. Rather than a synchronous landscape-wide outbreak, the results indicate a spatially modular propagation process shaped by semi-connected habitat patches, ecological corridors, and heterogeneous local amplification zones.

**Figure 4:**
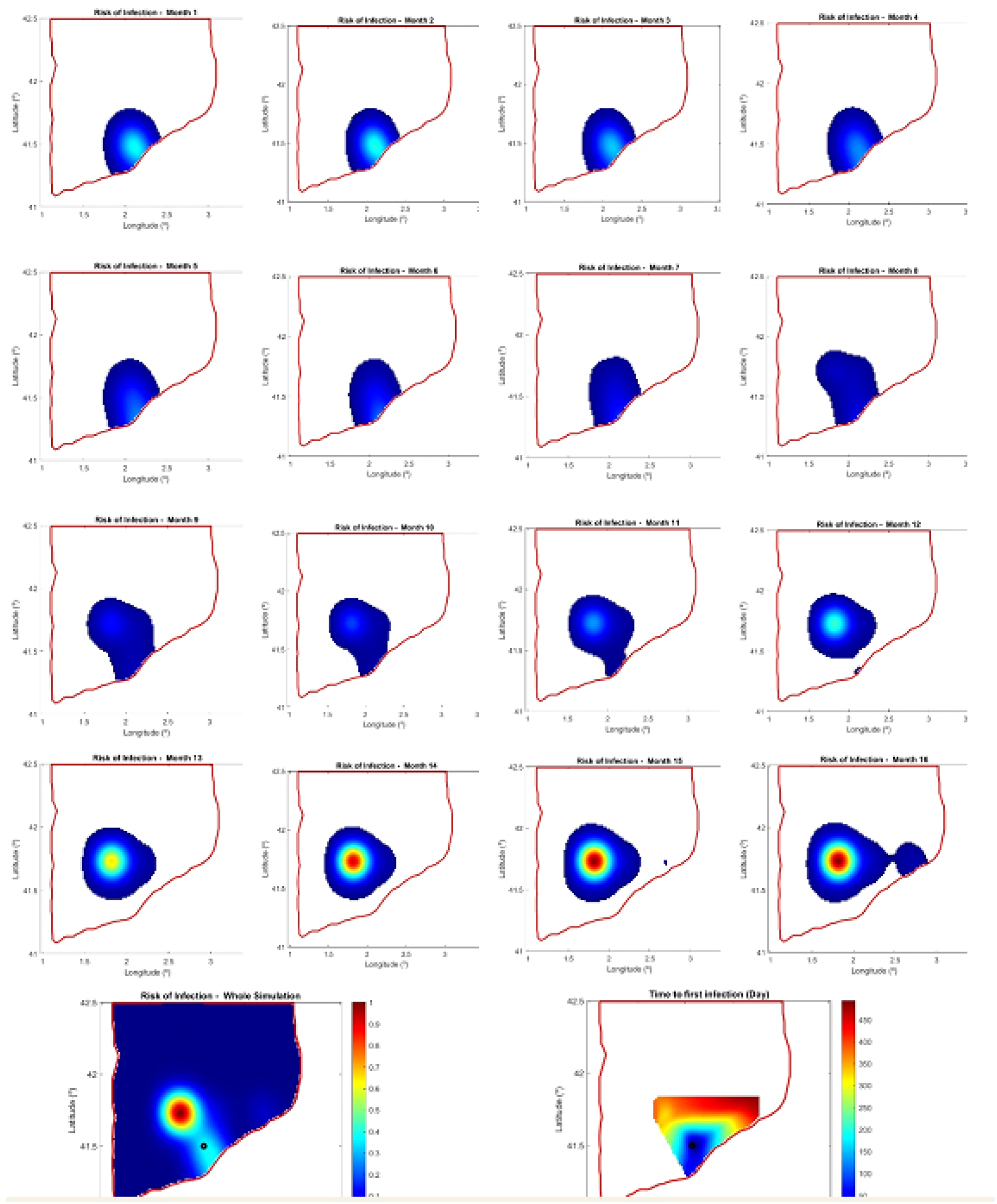
Monthly infection-risk dynamics of African swine fever (ASF) spread in wild boar populations under the Catalonia scenario. Sequence of spatial risk maps showing the normalised monthly infection probability relative to the maximum risk value observed across the full simulation scenario. The maps illustrate the progressive eco-epidemiological wavefront of disease spread during the 500-day modelling period, highlighting shifting spatial amplification zones, the delayed emergence of visible infection fronts, and temporally structur_1_e_9_d surveillance windows. Black dot represent the first case of ASF in wild boar in Catalonia.

This pattern is consistent with the ecology of peri-urban Mediterranean landscapes such as the Barcelona urban–natural gradient. In such systems, wild boar populations adapted to human-modified environments often show altered movement routines, flexible habitat use, and restructured contact networks driven by anthropogenic resources. Predictable waste-derived food sources, urban green infrastructure, agroforestry areas, and urban–natural refugia may generate behaviourally buffered mobility and heterogeneous aggregation patterns (Amici et al., 2012; Bosch et al., 2017a; Aguilar-Vega et al., 2024; Pérez-González et al., 2025). Such buffering may reduce synchronised mass-contact events while favouring stepwise dispersal and prolonged epidemiological persistence. In the affected Barcelona–Collserola–Vallès area, wild boar function as urban-adapted populations rather than strictly wild populations. Their ecology is shaped by a connected urban–natural interface, where behavioural plasticity leads to modified movement patterns, opportunistic resource use and restructured contact networks, with direct consequences for ASF transmission dynamics (Bosch and Ivorra et al., 2026a; Bosch et al., 2026b).

At monthly resolution, the outbreak behaves as a moving eco-epidemiological wavefront rather than as a fixed infected area. Some areas act as early transmission bridges, others as delayed amplification zones, and others as residual persistence pockets after the main front has advanced. This is operationally important because it defines shifting surveillance windows in space and time. Areas showing only moderate risk in one month may become epidemiologically critical shortly afterwards, particularly in fragmented peri-urban systems where detectability dynamics are complex. While carcasses may be more likely to be reported due to higher human presence, increased disturbance, indirect transmission opportunities and behavioural adaptation of wild boar can simultaneously complicate spatial containment and delay clear interpretation of the true infection front. Delayed carcass detection has been recognised as a major surveillance bias in wild boar ASF outbreaks, potentially masking the true spatial extent of infection during early epidemic phases (Probst et al., 2017; Ito & Bosch et al., 2024). Monthly risk surfaces therefore provide a basis for anticipatory surveillance, targeted carcass search, and adaptive deployment of control measures along likely acceleration corridors.

### 4.4. Field support and strain-dependent interpretation

Field evaluation and consistency with independent connectivity-based approaches and other studies further reinforce the eco-epidemiological interpretation of the WIM-BOARD outputs. Early field observations provide support for these modelled dynamics. In particular, the early southward phase closely matches the observed field evolution during the first weeks of the outbreak, including positive detections towards Sant Cugat del Vallès and the southern Collserola sector. This agreement reinforces the operational credibility of the analysis and suggests that the southern signal should not necessarily be interpreted as an anomalous jump, but rather as a plausible early dispersal phase following ecologically favourable routes within a fragmented yet still functionally connected urban–natural system.

This interpretation is further supported by previous connectivity analyses for the same region. Least-cost path modelling and functional landscape-connectivity assessments have shown that the outbreak area is structured by habitat patches linked through epidemiologically relevant corridors, including riparian strips, agroforestry mosaics, peri-urban vegetation belts, and residual connective elements between infrastructure barriers (Goicolea et al., 2024; Faustini et al., 2025). These spatial patterns are also supported by a broader European connectivity framework, for which harmonised cartographic datasets and spatial outputs are publicly available for consultation and download (Connectivity Modelling Project, 2024). This is in line with a broader body of work showing that wild boar movement and ASF spread are structured by non-Euclidean, habitat-dependent pathways rather than by simple geographic distance (Morelle and Lejeune, 2015; Podgórski et al., 2018). In particular, the route connecting the initial outbreak area near the UAB/Bellaterra sector with the southernmost positive detection runs close to a previously identified functional corridor. This corridor crosses riparian vegetation, agroforestry patches, and infrastructure interfaces, including the AP-7 highway area, suggesting that movement may be channelled through residual permeability rather than entirely prevented by formal barriers, as illustrated in Fig. 5, derived from European-scale ASF connectivity and ecological corridor mapping (Goicolea et al., 2024). This has direct implications for interpreting the southernmost positive live wild boar detected several kilometres from the initial cluster. In a fragmented peri-urban system such as Barcelona–Vallès, such a detection does not necessarily imply an abrupt epidemiological leap. More plausibly, it represents the detectable edge of an infection process progressing through ecologically functional space before becoming visible to surveillance. This interpretation is supported by the movement ecology of wild boar in these systems, which is non-linear, patch-dependent and strongly mediated by cover, resource availability, disturbance and social structure. A straight-line distance of 5–6 km is therefore compatible with a longer, tortuous movement pathway across mosaics of natural habitat, riparian corridors, agroforestry patches and linear infrastructures.

**Figure 5:**
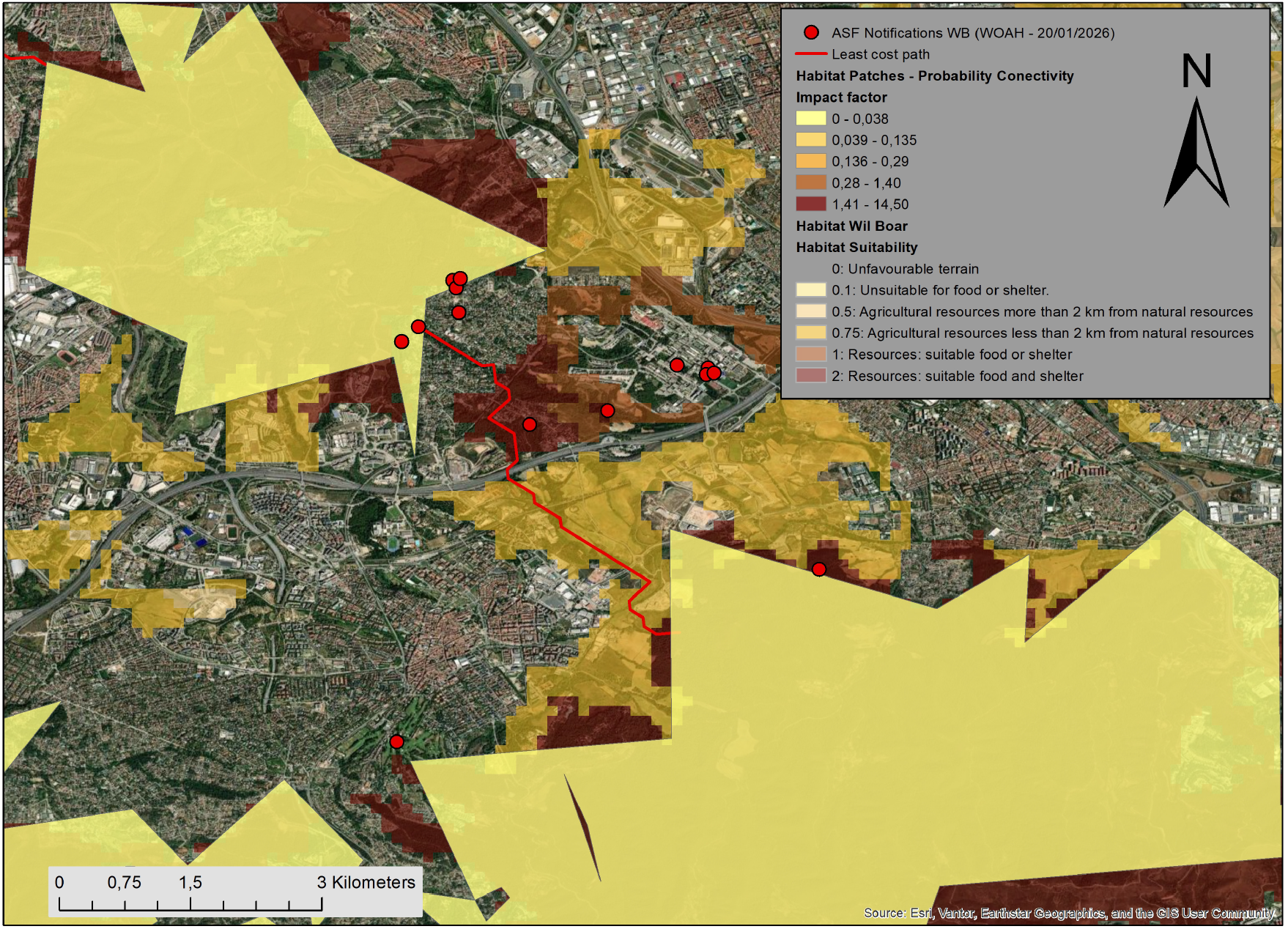
Least-cost path or functional corridor (red line) connecting the initial outbreak area near the UAB/Bellaterra sector with the southernmost ASF-positive detection prior to 20 January 2026. Habitat patch connectivity is represented in yellow as the impact factor of probability connectivity, while wild boar ASF notifications as of 20 January 2026 are shown in red. Habitat suitability is displayed as a semi-transparent gradient from light to dark brown, with darker areas indicating higher habitat quality (Bosch et al., 2017a). The corridor illustrates a plausible movement pathway through a fragmented peri-urban landscape structured by ecological connectivity and residual permeability across infrastructure barriers. Data derived from European-scale ASF connectivity and ecological corridor mapping (Goicolea et al., 2024).

The ecology of the Barcelona–Collserola–Vallès wild boar system further strengthens this interpretation. Wild boar in this setting occupy a hybrid ecological gradient in which urban areas function mainly as feeding grounds, while nearby natural patches provide refuge and reproductive space. Habitat modelling based on Quality of Available Habitat has shown that agro-urban and agroforestry resources located within approximately 2 km of natural habitat edges significantly increase the probability of recurrent wild boar use (Bosch et al., 2017a). Thus, the southern positive case should not be interpreted as urban spread detached from landscape ecology, but rather as spread rooted in a connected urban–natural wild boar system. Hydro-ecological features may also reinforce this pattern, as water-associated habitats, temporary stream systems, and low-resistance valley-bottom routes may facilitate both movement and environmental persistence (Ito & Bosch et al., 2024). Spatial network analysis of the hydrological system (rivers and streams) is illustrated in Fig. 6, where ASF notifications, least-cost pathways, habitat-connectivity patches, and the local river–stream network show a marked spatial convergence in the Barcelona–Vallès system, based on the hydrological risk framework described by Ito & Bosch et al. (2024).

**Figure 6:**
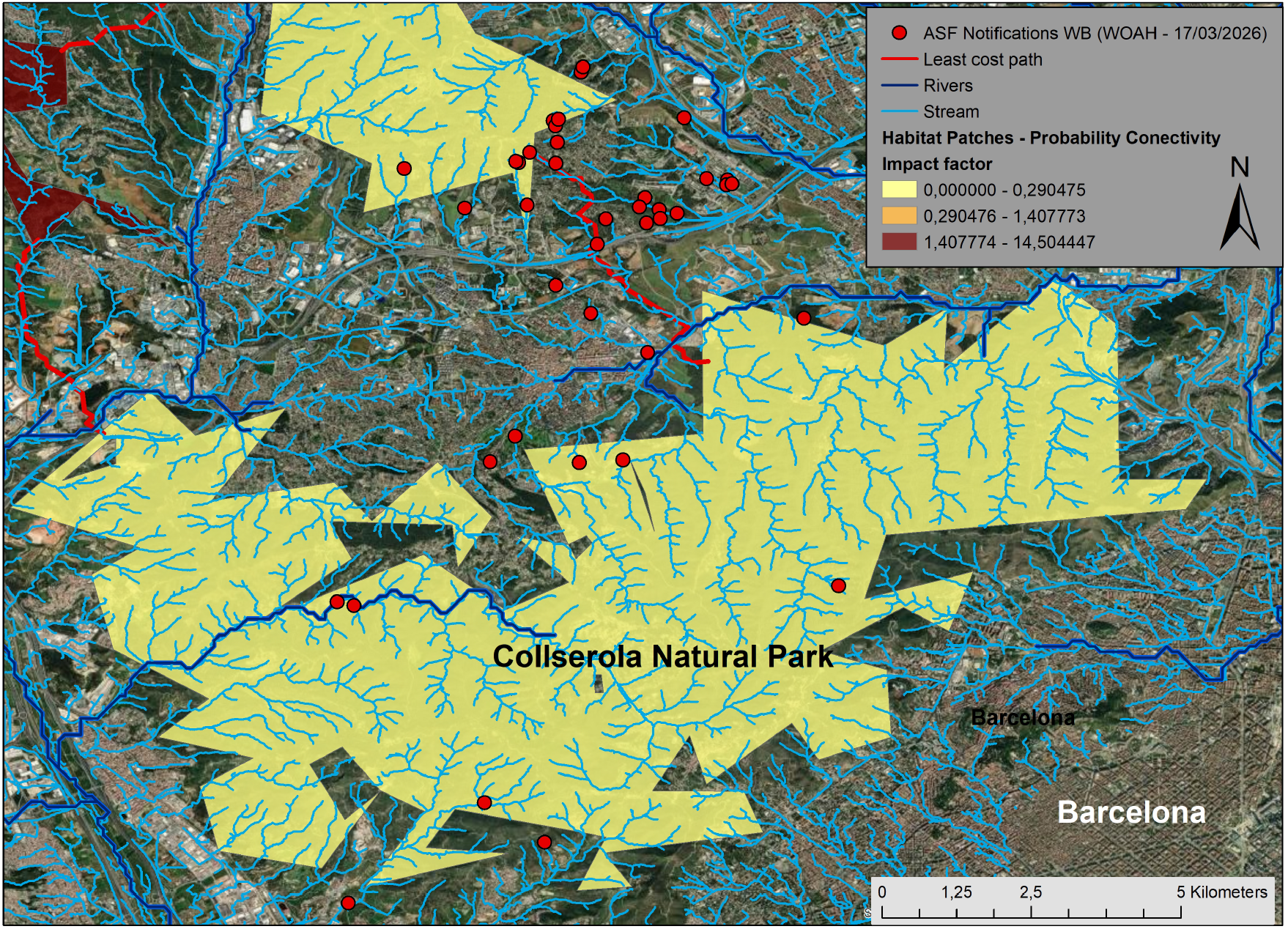
Hydro-ecological context of ASF notifications and functional connectivity in the Barcelona–Vallès system, based on the hydrological-risk framework described by Ito & Bosch et al. (2024). Light blue lines represent streams, whereas dark blue lines represent rivers with permanent water flow. Red points indicate ASF notifications in wild boar as of 17 March 2026, and the red line shows the least-cost path or functional corridor. Yellow to dark brown polygons represent habitat-patch probability connectivity and its associated impact factor. The satellite background highlights the AP-7 motorway, populated nuclei, and the fragmented habitat structure of the peri-urban landscape. The Collserola Natural Park and the city of Barcelona are located in the lower part of the figure. Together, these elements illustrate the spatial convergence between hydrological structure, ecological connectivity, and reported ASF cases in a fragmented urban–natural interface.

Hydrologically structured corridors may become even more relevant during the wet season, when temporary stream networks regain functional continuity and valley-bottom routes offer reduced movement resistance together with increased environmental suitability for virus persistence. In the Barcelona–Vallès system, least-cost connectivity pathways appear to converge with topographically low and erosion-prone zones beneath major transport infrastructures, such as motorway underpasses, where runoff concentration and geomorphological permeability may facilitate both wildlife passage and localised infection persistence. When formally recognised wildlife crossings are closed or disturbed, wild boar movements may become increasingly channelled through these residual hydrological or infrastructure-associated corridors, including railway margins and drainage-related erosion gaps, thereby intensifying directional transmission pathways across fragmented peri-urban landscapes. Importantly, this pattern does not imply waterborne transmission of ASF virus; rather, it reflects the role of hydrological networks as spatial structuring elements of wild boar movement ecology, where animals preferentially use valley-bottom environments for thermoregulation, resource access and movement, particularly under disease conditions, thereby increasing contact rates and environmental exposure along these corridors (Ito et al., 2024; Morelle et al., 2019b).

At the same time, the results must be interpreted in light of possible strain-dependent effects. In its current operational configuration, the modelling approach is parameterised primarily for highly virulent ASF strains, which is consistent with the simulated epidemic structure shown in Fig. 1. However, the comparatively moderate early observable mortality signal in the field, together with reports of seropositive animals and infected wild boar still mobile at the time of capture or selective removal, suggests that the detected virus may not behave exactly like a classical highly virulent genotype II strain. If infectious animals survive longer, they may disperse further, remain infectious for longer periods, and generate more diffuse transmission before detection. In this sense, the present outputs should be interpreted as a conservative baseline scenario under acute-strain assumptions, while acknowledging that future incorporation of attenuated-strain dynamics (Martínez & Bosch et al., 2023) may broaden the range of plausible epidemiological scenarios.

### 4.5. Integrated interpretation and operational implications

Taken together, these results support the interpretation of the Catalonia outbreak as a landscape-mediated epidemiological process shaped by functional connectivity, wildlife behavioural adaptation, peri-urban ecological structure, and delayed detectability. This interpretation is consistent with previous large-scale analyses across Eastern Europe, which have shown that ASF spread in wild boar populations is strongly structured by landscape-level dispersal ecology rather than by simple homogeneous diffusion processes (Podgórski and Śmietanka, 2018). Similar landscape-structured spread dynamics have also been reported in Central Europe, where epidemiological expansion in wild boar populations followed spatially constrained ecological pathways (Bosch et al., 2017a,b; Sauter-Louis et al., 2020; Goicolea et al., 2024). Viewed from this integrative perspective, rather than as a set of isolated outputs, the combination of temporal trajectories, cumulative risk patterns, directional dispersal phases, monthly propagation surfaces, and field-supported connectivity evidence provides a coherent eco-epidemiological understanding of ASF emergence at a complex wildlife–urban interface. This interpretative capacity arises from the inherently multidisciplinary architecture of the modelling framework, which integrates applied mathematics, epidemiology, wildlife ecology, veterinary science, environmental processes, and management-oriented knowledge within a unified analytical environment. Such interdisciplinary integration is essential for translating complex spatial epidemiological dynamics into biologically grounded and operationally actionable decision-support outputs.

From a management perspective, the applicability of these findings is direct. Surveillance should be intensified not only within the initial outbreak cluster but also along ecologically plausible directions of spread, while carcass search should prioritise functionally connected corridors, riparian belts and residual permeability points. Fencing strategies should focus on likely crossing areas rather than assumed barrier lines, and population-management interventions should be timed before spring reproductive amplification (Bosch et al., 2020b), particularly under scenarios in which virus establishment in wild boar remains a risk. Operational guidelines for ASF management similarly emphasise the importance of spatially targeted population reduction and carcass removal adapted to landscape context (Guberti et al., 2019). In highly anthropised systems, these measures must be complemented by appropriate waste management that reduces food accessibility for wildlife, reinforced public communication, access control, and strict attention to indirect human-mediated transmission pathways(Aguilar-Vega et al., 2024). More specifically, the results suggest that management restricted to the urban core is unlikely to be sufficient, as wild boar populations remain functionally connected to neighbouring municipalities. Effective control of population abundance therefore requires coordinated strategies across urban, peri-urban and surrounding natural areas.

In addition, WIMBOARD provides a quantitative framework for designing and optimising wildlife population-management strategies under ASF scenarios. Wild boar populations in Europe have experienced sustained demographic expansion driven by anthropogenic landscape changes, increased food availability and reduced predation pressure (Barrios-García and Ballari, 2012; Bosch et al., 2012; Amici et al., 2012; Tack, 2018; Aguilar-Vega et al., 2024), resulting in high-density populations that can facilitate pathogen persistence and spatial spread. Within this context, population reduction has been widely considered as a complementary control measure, although its effectiveness depends critically on spatial targeting, timing and ecological context (Keuling et al., 2013; Pepin et al., 2020).

Through scenario-based simulations, WIMBOARD enables estimation of where, when and to what extent wild boar populations should be reduced in order to disrupt transmission pathways and limit epidemic expansion. By integrating host movement, functional connectivity and infection dynamics, the platform identifies priority intervention zones along predicted spread corridors and quantifies the intensity of selective removal strategies, including controlled hunting and trapping. This supports the design of spatially structured population-reduction schemes aimed at reducing epidemiological connectivity, lowering contact rates and establishing functional low-density buffer zones between infected areas and disease-free regions. Such forward-looking simulations provide an operational tool for adaptive management, supporting the transition from reactive population control towards anticipatory, evidence-based strategies aligned with predicted outbreak trajectories.

While ASF spread in wild boar populations represents the primary ecological driver of landscape-scale transmission dynamics, the ultimate sanitary and socio-economic impact of outbreaks is largely determined by the risk of spillover into domestic pig production systems and associated human activities. Furthermore, the spatially explicit risk outputs may also support assessment of potential exposure at the wildlife–livestock interface, particularly in areas where functional landscape connectivity may facilitate indirect transmission processes. Farm biosecurity should therefore be interpreted not only as an on-farm measure but also as dependent on broader landscape-environment and ecological interface conditions. This perspective is epidemiologically relevant because both direct and indirect interactions between infected wild boar populations and domestic pig production systems may influence spillover risk (Cadenas-Fernández et al., 2019; Bosch et al., 2020a). Although farm biosecurity remains a cornerstone of ASF prevention, increasing evidence indicates that virus introduction probability is also modulated by landscape-scale infection pressure, environmental contamination and wildlife connectivity patterns (Costard et al., 2013; Chenais et al., 2018). Even holdings with high internal biosecurity standards may experience elevated exposure risk when located within functionally connected areas with sustained wild boar infection circulation. Under such conditions, indirect transmission pathways mediated by contaminated environments, mechanical vectors or episodic wildlife–livestock interface contacts may contribute to delayed spillover events despite the implementation of recommended preventive measures (Mur et al., 2012). In addition, human-mediated factors — including farmer behaviour, personnel routines, visitor movements and operational habits at the wildlife–farm interface — may further modulate transmission probability. From an eco-epidemiological perspective, landscape-mediated epidemiological pressure operating through indirect wildlife–environment–human interfaces may elevate and delay context-dependent spillover risk beyond strictly on-farm biosecurity conditions or barriers. Because farm-level geolocation data are sensitive and no specific farm-level analysis is presented here, this aspect is discussed only as a potential applied extension of the modelling framework. Such operational analyses are designed to support decision-making in emerging outbreak contexts, particularly in situations where real-time data availability is limited. These analyses are available and may be shared upon request with competent authorities and stakeholders, with the aim of supporting context-specific preparedness exercises and informing strategic risk mitigation planning.

Beyond these immediate operational considerations, the modelling framework also allows exploration of longer-term epidemiological trajectories under baseline intervention scenarios. In particular, long-term exploratory simulations extending the temporal horizon up to 1,000 days (approximately 2.7 years) were conducted under baseline assumptions reflecting currently implemented control measures in order to explore potential medium- to long-term ecological spread trajectories. These forward-looking analyses were designed to examine plausible future expansion directions, the progressive emergence of risk in new municipalities, potential exposure of natural ecosystems and protected areas, and the increasing environmental interface risk for pig-production systems located within functionally connected landscapes. Such projections must be interpreted cautiously, as wildlife disease dynamics are inherently influenced by stochastic dispersal processes, behavioural adaptation, and evolving management responses, and they do not explicitly account for human-mediated transmission. This limitation is particularly relevant in highly anthropised landscapes, where indirect human-related mechanisms may substantially alter the progression, directionality, and apparent speed of the epidemic wave. This uncertainty is consistent with previous conceptual analyses suggesting that ASF spread in wild boar may follow either rapid epidemic waves or slower persistence-driven dynamics depending on ecological context and strain characteristics (Schulz et al., 2019). Nevertheless, these simulations provide a valuable anticipatory perspective for preparedness planning by illustrating how infection could continue to propagate under a baseline intervention scenario, thereby helping translate predictive modelling outputs into structured support for decision-making under uncertainty. Although not included within the core analytical results because of their exploratory nature, these extended projections are operationally available upon request to competent authorities and stakeholders to support context-specific preparedness exercises and strategic risk-mitigation planning.

Although soft ticks of the *Ornithodoros erraticus* complex played an important role as biological vectors during the historical ASF epidemic in parts of the Iberian Peninsula (Boinas et al., 2011; Sánchez-Vizcaíno et al., 2012), their relevance in the current Catalonia outbreak appears negligible. Based on the information currently available, the presence of this vector has not been detected in the affected area (Boinas et al., 2014; GBIF, 2023). Accordingly, tick-mediated transmission is not considered a relevant component of the present outbreak scenario, although it remains a historical reminder that ASF persistence mechanisms may vary across ecological contexts.

Experimental infection studies in wild boar have provided quantitatively grounded parameters that substantially refine the interpretation of ASF transmission dynamics under variable virulence conditions (Martínez & Bosch et al., 2023). In particular, empirical parameterisation of attenuated genotype II infection scenarios has yielded an experimental intragroup reproduction value (*R_x_*) of approximately 4.5 and demonstrated that infection outcomes may include prolonged infectiousness, heterogeneous mortality trajectories, and persistent epidemiological tails extending well beyond the acute outbreak phase. Under mixed-virulence assumptions, studies have shown that between 4% and 17% of individuals may remain intermittently or permanently infectious after the epidemic peak, while an additional 5% to 11% of animals may survive infection in a recovered-but-susceptible state, retaining the potential for re-infection upon exposure to highly virulent strains. In some experimental modelling configurations, persistence of infectiousness exceeded two years, highlighting a substantially longer epidemiological tail than would be expected under classical acute-strain transmission frameworks. These biologically grounded findings are particularly relevant for interpreting the Catalonia outbreak, where field observations of seropositive wild boar, delayed carcass detection, and variable mortality timing suggest the possibility of heterogeneous post-infection states influencing spatial transmission patterns. Integrating such experimentally derived parameters into predictive modelling frameworks enhances the ecological realism of outbreak interpretation and supports the exploration of broader virulence-spectrum scenarios relevant for surveillance prioritisation and containment planning. These scenario-based projections support the improvement of surveillance strategies and early detection systems by translating modelling outputs into actionable decision-support tools under conditions of epidemiological uncertainty.

Table 1 summarises the main conceptual links between modelling outputs, wildlife behavioural ecology, landscape structure, and their operational implications under the 2025–2026 Catalonia outbreak scenario.

**Table 1:** Conceptual eco-epidemiological interpretation linking modelling outputs, wildlife behavioural ecology, and landscape structure under the Catalonia ASF scenario.

| Component | Landscape-ecological mechanism | Expected effect | Operational implication |
| --- | --- | --- | --- |
| Landscape fragmentation | Reduced habitat continuity and compartmentalisation of wild boar subpopulations | Progressive rather than explosive transmission dynamics | Spatially differentiated surveillance buffers |
| Wildlife urban adaptation | Modified movement routines and flexible resource use in human-dominated environments | Altered contact networks and asynchronous infection spread in urban–natural interface settings | Surveillance of peri-urban interfaces and waste-related hotspots |
| Anthropogenic food resources | Aggregation around predictable human-derived food sources | Localised persistence and recurrent transmission foci | Targeted carcass search and communication in human-modified areas |
| Corridor-mediated connectivity | Functional ecological links despite infrastructure barriers | Directional and wave-like epidemic expansion | Strategic placement of fencing, traps, and population-management actions |
| Behavioural buffering and modular mobility | Stepwise dispersal between semi-connected habitat patches | Lower early observable mortality, prolonged epidemiological pressure, and structured spatial diffusion | Anticipatory surveillance based on predicted spread sequence |
| Targeted population reduction | Spatially structured reduction of host abundance along predicted transmission corridors | Reduced epidemiological connectivity, contact rates, and directional spread | Scenario-based definition of the intensity, timing, and targeting of selective removal needed to create functional disease-buffer zones |
| Integrated modelling interpretation | Coupling temporal prevalence dynamics with cumulative spatial risk surfaces | Landscape-mediated epidemiological modulation | Use of the modelling approach as an operational decision-support tool |
| Long-term scenario exploration | Progressive redistribution of epidemiological pressure across connected habitats over extended horizons | Delayed emergence of secondary risk zones and shifting surveillance priorities | Extended-horizon simulations for preparedness planning and adaptive control design |
| Wildlife–livestock interface risk | Overlap between predicted wild boar infection corridors and pig-production environments | Landscape-mediated exposure risk and indirect and delayed spillover pathways | Prioritisation of farm biosecurity, targeted surveillance, and contingency planning |

## 5. Conclusions

The 2025–2026 Catalonia outbreak shows that African swine fever (ASF) spread in wild boar populations within highly anthropised peri-urban environments cannot be adequately understood through simplified assumptions of homogeneous diffusion or purely reactive outbreak interpretation. The analyses presented here support the interpretation of ASF spread in this setting as a landscape-mediated epidemiological process shaped by functional connectivity, host behavioural adaptation, fragmented urban–agro–natural mosaics, and delayed detectability.

The results indicate that outbreak expansion was structured and directional rather than isotropic, and that visible mortality may substantially lag behind the true spatial extent of viral circulation. In particular, the identification of an early southward signal consistent with field observations and of a north–northwest expansion scenario with increased potential for regional amplification and persistence highlights the importance of interpreting ASF dynamics through the combined lens of landscape connectivity and peri-urban wildlife ecology. These findings have direct implications for surveillance and control, as carcass-based passive surveillance may underestimate the real extent of infection, whereas cumulative risk patterns and monthly propagation surfaces can help identify anticipatory surveillance windows and ecologically plausible directions of spread.

More broadly, this study illustrates the value of spatially explicit eco-epidemiological modelling for supporting outbreak interpretation, surveillance prioritisation, and preparedness planning under complex field conditions. At the same time, the present analysis has important limitations. First, the current modelling configuration is parameterised primarily for acute high-virulence ASF scenarios and does not yet incorporate attenuated-strain dynamics, although such a module is under development. Second, the study is predictive in nature and explores plausible spread trajectories over a 500-day horizon on the basis of information available during the early stage of the outbreak; consequently, longer-term projections should be interpreted with caution. In addition, the relative importance of spatial expansion patterns identified in this study should be understood within the temporal limits of the 500-day simulation horizon, as their epidemiological relevance may evolve under longer-term dynamics.

Ultimately, the ecological dynamics of ASF in wild boar become operationally relevant when landscape-mediated epidemiological pressure translates into context-dependent spillover risk for domestic pig production systems. Functional connectivity and indirect wildlife–environment–human interface processes may sustain infection pressure beyond farm-level biosecurity barriers, potentially contributing to delayed spillover events.

Designed for continental-scale epidemiological modelling, WIMBOARD generates fine-scale spatial intelligence to guide local surveillance and outbreak control decisions. The Catalonia case study illustrates how the platform can provide near real-time projections to support proactive, stakeholder-oriented decision-making under uncertainty, effectively shifting outbreak management from reactive to anticipatory frameworks.

Taken together, these results provide a biologically grounded and operationally relevant framework for interpreting ASF spread in complex peri-urban wild boar systems and for supporting more proactive surveillance and containment strategies under conditions of epidemiological uncertainty.

## Acknowledgements

This research was supported by the Horizon 2020 programme of the European Union through the project *VACDIVA – A Safe DIVA vaccine for African Swine Fever control and eradication* (Grant Agreement No. 862874; DOI: 10.3030/862874). This work was also supported by project PID2023-146754NB-I00, funded by MCIU/AEI/10.13039/501100011033 and by the European Regional Development Fund (FEDER, EU), and by project PCI2024-153478 under the European M-ERA.Net programme, funded by MCIU/AEI/10.13039/501100011033 and the European Union.

## References

Aguilar-Vega, C., Sánchez-Vizcaíno, J.M., Bosch, J., 2024. Identifying sites where wild boars can consume anthropogenic food waste with implications for African swine fever. PLOS ONE 19, e0308502. doi:10.1371/journal.pone.0308502.

Alexander, N.S., Massei, G., Wint, W., 2016. The european distribution of Sus scrofa: model outputs from the project described within the poster “where are all the boars?” an attempt to gain a continental perspective. Open Health Data 4, e1. doi:10.5334/ohd.2.

Amici, A., Serrani, F., Rossi, C.M., Primi, R., 2012. Increase in crop damage caused by wild boar (Sus scrofa l.): the “refuge effect”. Agronomy for Sustainable Development 32, 683–692. URL: https://dspace.unitus.it/handle/2067/35075.

Arias, M., Sánchez-Vizcaíno, J.M., 2002. African swine fever eradication: The spanish model, in: Morilla, A., Yoon, K.J., Zimmerman, J.J. (Eds.), Trends in Emerging Viral Infections of Swine. Iowa State Press, Ames, pp. 133–139.

Barrios-García, M.N., Ballari, S.A., 2012. Impact of wild boar (Sus scrofa) in its introduced and native range: a review. Biological Invasions 14, 2283–2300. doi:10.1007/s10530-012-0229-6.

Blome, S., Franzke, K., Beer, M., 2020. African swine fever – a review of current knowledge. Virus Research 287, 198099. doi:10.1016/j.virusres.2020.198099.

Boinas, F., Ribeiro, R., Madeira, S., Palma, M., de Carvalho Ferreira, H.C., 2011. African swine fever in Ornithodoros erraticus ticks. Vector-Borne and Zoonotic Diseases 11, 1535–1540. doi:10.1089/vbz.2011.0666.

Boinas, F.S., Ribeiro, R., Madeira, S., 2014. The medical and veterinary role of ornithodoros erraticus complex ticks (acari: Ixodida) on the iberian peninsula. Journal of Vector Ecology 39, 238–248. doi:10.1111/jvec.12100.

Bosch, J., Barasona, J.A., Cadenas-Fernandez, E., Jurado, C., Pintore, A., Denurra, D., Vicente, J., Sánchez-Vizcaíno, J.M., 2020a. Retrospective spatial analysis for african swine fever in endemic areas to assess interactions between susceptible host populations. PLOS ONE 15, e0233473. doi:10.1371/journal.pone.0233473.

Bosch, J., Iglesias, I., Martínez, M., de la Torre, A., 2020b. Climatic and topographic tolerance limits of wild boar in eurasia: implications for their expansion. Geography, Environment, Sustainability 13, 107–114. doi:10.24057/2071-9388-2019-52.

Bosch, J., Iglesias, I., Muñoz, M.J., De la Torre, A., 2017a. A cartographic tool for managing African swine fever in Eurasia: mapping wild boar distribution based on the quality of available habitats. Transboundary and Emerging Diseases 64, 1720–1733. doi:10.1111/tbed.12559.

Bosch, J., Ivorra, B., Aguilar-Vega, C., Ito, S., Sánchez-Vizcaíno, J.M., Ramos, Á.M., 2026a. WIMBOARD: An operational eco-epidemiological decision-support platform. application to the 2025–2026 African swine fever outbreak in wild boar in catalonia. URL: https://www.researchgate.net/publication/402537835_WIMBOARD_An_Operational_Eco-Epidemiological_Decision-Support_Platform_Application_to_the_2025-26_African_Swine_Fever_Outbreak_in_Wild_Boar_in_Catalonia. preprint, March 2026.

Bosch, J., Ivorra, B.P.P., Aguilar-Vega, C., Ito, S., Sánchez-Vizcaíno Rodríguez, J.M., Ramos del Olmo, Á.M., 2026b. African Swine Fever in Spain (Catalonia, November 2025): Lessons Learned from Science, Surveillance, and Field Response. URL: https://docta.ucm.es/entities/publication/4a994792-9c99-4264-abb4-960382b7dc3d.

Bosch, J., Peris, S., Fonseca, C., Martinez, M., De la Torre, A., Iglesias, I., Muñoz, M.J., 2012. Distribution, abundance and density of the wild boar on the Iberian Peninsula, based on the CORINE program and hunting statistics. Folia Zoologica 61, 138–151. doi:10.25225/fozo.v61.i2.a7.2012.

Bosch, J., Rodríguez, A., Iglesias, I., Muñoz, M.J., Jurado, C., Sánchez-Vizcaíno, J.M., De la Torre, A., 2017b. Update on the risk of introduction of African swine fever by wild boar into disease-free European Union countries. Transboundary and Emerging Diseases 64, 1424–1432. doi:10.1111/tbed.12527.

Cadenas-Fernández, E., Sánchez-Vizcaíno, J.M., Pintore, A., Denurra, D., Jurado, C., Vicente, J., Barasona, J.A., 2019. Free-ranging wild boar and domestic pig interactions: implications for disease transmission. Frontiers in Veterinary Science 6, 154. doi:10.3389/fvets.2019.00154.

Chenais, E., Ståhl, K., Guberti, V., Depner, K., 2018. Identification of wild boar–habitat epidemiologic cycle in African swine fever epizootic. Emerging Infectious Diseases 24, 810–812. doi:10.3201/eid2404.172127.

Connectivity Modelling Project, 2024. Asf wild boar connectivity and corridors dataset. https://docs.google.com/forms/d/e/1FAIpQLSeGd8xq2L_2ZH47aVR4lryFYzKbL5rEHuv6NAZE-wrkIltxtg/viewform. Cartography download portal.

Costard, S., Mur, L., Lubroth, J., Sánchez-Vizcaíno, J.M., Pfeiffer, D.U., 2013. Epidemiology of African swine fever virus. Virus Research 173, 191–197. doi:10.1016/j.virusres.2012.10.030.

Dixon, L.K., Stahl, K., Jori, F., Vial, L., Pfeiffer, D.U., 2020. African swine fever epidemiology and control. Annual Review of Animal Biosciences 8, 221–246. doi:10.1146/annurev-animal-021419-083741.

European Commission, 2020. Strategic approach to the management of african swine fever for the eu. URL: https://food.ec.europa.eu/system/files/2020-04/ad_control-measures_asf_wrk-doc-sante-2015-7113.pdf.

European Commission, 2023. African swine fever: prevention, control and eradication guidelines in the european union. URL: https://food.ec.europa.eu/animals/animal-diseases/diseases-and-control-measures/african-swine-fever_en.

European Commission, VACDIVA Consortium, 2024. VACDIVA: A safe DIVA vaccine for African swine fever control and eradication. doi:10.3030/862874. Horizon 2020 Project, Grant Agreement No. 862874.

Faustini, G., Soret, M., Defossez, A., Bosch, J., Conte, A., Tran, A., 2025. Habitat suitability mapping and landscape connectivity analysis to predict African swine fever spread in wild boar populations: a focus on Northern Italy. PLOS ONE 20, e0317577. doi:10.1371/journal.pone.0317577.

Fernández-Carrión, E., Ivorra, B., Martínez-López, B., Ramos, Á.M., Sánchez-Vizcaíno, J.M., 2016. Implementation and validation of an economic module in the Be-FAST model to predict costs generated by livestock disease epidemics: application to classical swine fever epidemics in Spain. Preventive Veterinary Medicine 126, 66–73. doi:10.1016/j.prevetmed.2016.01.015.

Gallardo, C., Soler, A., Rodze, I., Nieto, R., Cano-Gómez, C., Fernández-Pinero, J., Arias, M., 2015. African swine fever : a global view of the current challenge. Porcine Health Management 1, 21. doi:10.1186/s40813-015-0013-y.

Gámbaro, F., et al., 2025. Viral dna to explore african swine fever genotype ii in europe. Genome Biology and Evolution Estimated evolutionary rate ∼ 5.7 × 10^−6^ substitutions per site per year.

GBIF, 2023. Gbif occurrence download: Ornithodoros erraticus distribution. URL: https://www.gbif.org. accessed November 2025.

Gervasi, V., et al., 2022. Combining hunting and intensive carcass removal to control african swine fever in wild boar populations. Preventive Veterinary Medicine 201, 105594.

Goicolea, T., Cisneros-Araújo, P., Vega, C.A., Sánchez-Vizcaíno, J.M., Mateo-Sánchez, M., Bosch, J., 2024. Landscape connectivity for predicting the spread of ASF in the European wild boar population. Scientific Reports 14, 3414. doi:10.1038/s41598-024-53869-5.

Guberti, V., Khomenko, S., Masiulis, M., Kerba, S., 2019. African swine fever in wild boar: ecology and biosecurity. FAO Animal Production and Health Manual, FAO, Rome. URL: https://openknowledge.fao.org/server/api/core/bitstreams/de800dfa-d892-4238-bbad-854840a89ddc/content.

Guinat, C., Gogin, A., Blome, S., Keil, G., Pollin, R., Pfeiffer, D.U., Dixon, L., 2016. Transmission routes of african swine fever virus to domestic pigs: current knowledge and future research directions. Veterinary Record 178, 262–267. doi:10.1136/vr.103593.

Ito, S., Bosch, J., Aguilar-Vega, C., Jeong, H., Sánchez-Vizcaíno, J.M., 2024. Geospatial analysis for strategic wildlife disease surveillance: African swine fever in South Korea (2019–2021). PLOS ONE 19, e0305702. doi:10.1371/journal.pone.030 5702.

Ivorra, B., Martínez-López, B., Sánchez-Vizcaíno, J.M., Ramos, Á.M., 2014. Mathematical formulation and validation of the Be-FAST model for Classical Swine Fever Virus spread between and within farms. Annals of Operations Research 219, 25–47. doi:10.1007/s10479-012-1257-4.

Keuling, O., Baubet, E., Duscher, A., Ebert, C., Fischer, C., Monaco, A., Podgórski, T., Prévot, C., Ronnenberg, K., Sodeikat, G., Stier, N., Thurfjell, H., 2013. Mortality rates of wild boar Sus scrofa in europe. European Journal of Wildlife Research 59, 805–814. doi:10.1007/s10344-013-0733-1.

Lange, M., Thulke, H.H., 2017. Simulation modelling to support African swine fever control strategies. Scientific Reports 7, 1–12. doi:10.1038/s41598-017-09229-5.

Martínez, M.A., Bosch, J., Ivorra, B., Ramos, Á.M., Ito, S., Barasona, J.Á., Sánchez-Vizcaíno, J.M., 2023. Epidemiological impacts of attenuated African swine fever virus circulating in wild boar populations. Research in Veterinary Science 162, 104964. doi:10.1016/j.rvsc.2023.104964.

Martínez-López, B., Ivorra, B., Ngom, D., Ramos, Á.M., Sánchez-Vizcaíno, J.M., 2012. A novel spatial and stochastic model to evaluate the within and between farm transmission of classical swine fever virus: II. validation of the model. Veterinary Microbiology 155, 21–32. doi:10.1016/j.vetmic.2011.08.008.

Martínez-López, B., Ivorra, B., Ramos, Á.M., Sánchez-Vizcaíno, J.M., 2011. A novel spatial and stochastic model to evaluate the within and between farm transmission of classical swine fever virus: I. general concepts and description of the model. Veterinary Microbiology 147, 300–309. doi:10.1016/j.vetmic.2010.07.009.

Morelle, K., Bubnicki, J., Churski, M., Gryz, J., Podgórski, T., Kuijper, D.P.J., 2019a. Disease-induced mortality outweighs hunting in shaping wild boar population dynamics. Ecological Applications 29, e01865. doi:10.1002/eap.1865.

Morelle, K., Jezek, M., Licoppe, A., Podgórski, T., 2019b. Deathbed choice by asf-infected wild boar can help find carcasses. Transboundary and Emerging Diseases 66, 1821–1826. doi:10.1111/tbed.13267.

Morelle, K., Lejeune, P., 2015. Seasonal variations of wild boar sus scrofa distribution in agricultural landscapes: a species distribution modelling approach. European Journal of Wildlife Research 61, 45–56. doi:10.1007/s10344-014-0872-6.

Mur, L., Martinez-Lopez, B., Sanchez-Vizcaino, J.M., 2012. Risk of african swine fever introduction into the eu through transport-associated routes. Transboundary and Emerging Diseases 59, 301–312.

Nefedeva, M., et al., 2020. Recombination shapes african swine fever virus serotype diversity. Scientific Reports DNA viruses show lower mutation rates than RNA viruses.

Pepin, K.M., Golnar, A.J., Abdo, Z., Podgórski, T., 2020. Ecological drivers of African swine fever virus persistence in wild boar populations. Transboundary and Emerging Diseases 67, 1616–1626. doi:10.1111/tbed.13426.

Pérez-González, J., Hidalgo-Toledo, S.P., Martínez, R., Hermoso-de Mendoza, J., Gonçalves, P., Hidalgo-de Trucios, S.J., 2025. Demography, peri-urban presence, and male-biased dispersal in an expanding wild boar (Sus scrofa) population in western Spain. European Journal of Wildlife Research 71, 149. doi:10.1007/s10344-025-02030-2.

Pittiglio, C., Khomenko, S., Beltran-Alcrudo, D., 2018. Wild boar mapping using population-density statistics: from polygons to high-resolution raster maps. PLOS ONE 13, e0193295. doi:10.1371/journal.pone.0193295.

Podgórski, T., Śmietanka, K., 2018. Do wild boar movements drive the spread of African swine fever? Transboundary and Emerging Diseases 65, 1588–1596. doi:10.1111/tbed.12910.

Podgórski, T., Apollonio, M., Keuling, O., 2018. Contact rates in wild boar populations: implications for disease transmission. Journal of Wildlife Management 82, 1210–1218. doi:10.1002/jwmg.21480.

Probst, C., Globig, A., Knoll, B., Conraths, F.J., Depner, K., 2017. Behaviour of wild boar carcasses and implications for African swine fever surveillance. Trans-boundary and Emerging Diseases 64, 205–212. doi:10.1111/tbed.12391.

Sánchez-Vizcaíno, J.M., Mur, L., Gomez-Villamandos, J.C., Carrasco, L., 2015. An update on the epidemiology and pathology of African swine fever. Journal of Comparative Pathology 152, 9–21. doi:10.1016/j.jcpa.2014.09.003.

Sánchez-Vizcaíno, J.M., Mur, L., Martínez-López, B., 2012. African swine fever: an epidemiological update. Transboundary and Emerging Diseases 59, 27–35. doi:10.1111/j.1865-1682.2011.01293.x.

